# Cooperativity and Avidity in Membrane Binding by C2AB Tandem Domains of Synaptotagmins 1 and 7

**DOI:** 10.1101/393702

**Authors:** H. Tran, L. Anderson, J. Knight

## Abstract

Synaptotagmin-1 (Syt-1) and synaptotagmin-7 (Syt-7) contain analogous tandem C2 domains, C2A and C2B, which together serve as a Ca^2+^ sensor to bind membranes and promote the stabilization of exocytotic fusion pores. Functionally, Syt-1 triggers fast release of neurotransmitters, while Syt-7 is involved in lower-Ca^2+^ processes such as hormone secretion. Evidence suggests that Syt-1 C2 domains bind membranes cooperatively, penetrating farther into membranes as the C2AB tandem than as individual C2 domains. In contrast, we previously reported that the two C2 domains of Syt-7 bind membranes independently, based in part on measurements of their liposome dissociation kinetics. Here, we have investigated the effects of C2A-C2B interdomain cooperativity with Syt-1 and Syt-7 using directly comparable measurements. We report Ca^2+^ sensitivities, dissociation kinetics, and membrane insertion using liposomes approximating physiological lipid compositions. Equilibrium Ca^2+^ titrations confirm that the Syt-7 C2AB tandem has a greater Ca^2+^ sensitivity of membrane binding than either of its individual domains. Stopped-flow fluorescence kinetic measurements show that Syt-1 C2AB dissociates from liposome membranes much more slowly than either of its isolated C2 domains, suggesting that the two C2 domains of Syt-1 bind membranes cooperatively. In contrast, the dominant population of Syt-7 C2AB has a dissociation rate comparable to its C2A domain, indicating a lack of cooperativity, while only a small subpopulation dissociates at a slower rate. Measurements using an environment-sensitive fluorescent probe indicate that the Syt-7 C2B domain inserts more deeply into membranes as part of the C2AB tandem, similarly to Syt-1. Overall, these measurements are consistent with a model in which the structural linkage of C2A and C2B impacts the membrane-binding geometry of synaptotagmin C2B domains, but imparts little or no cooperativity to Syt-7 membrane binding and dissociation events that are dominated by its C2A domain.

## Introduction

Synaptotagmins are a family of proteins that trigger and regulate fusion of secretory vesicles with the plasma membrane during exocytosis (1-3). Structurally, synaptotagmins contain an N-terminal transmembrane region, a cytoplasmic juxtamembrane linker region, and tandem C-terminal C2 domains, termed C2A and C2B. In eight of the seventeen mammalian isoforms, the C2 domains together function as a Ca^2+^ sensor to trigger SNARE-mediated membrane fusion (3-5). Although the synaptotagmins are structurally homologous, their Ca^2+^ and membrane binding sensitivities vary (6).

This study focuses on C2 domains from Syt-1 and Syt-7, two isoforms that serve as models for high-speed and high-sensitivity Ca^2+^-dependent vesicle fusion, respectively (7). Syt-1 serves as a primary Ca^2+^ sensor in the fast, synchronous release of neurotransmitters, while Syt-7 is involved in asynchronous neurotransmitter release, vesicle replenishment, synaptic facilitation, and endocrine and neuroendocrine secretion processes that occur at relatively low Ca^2+^ concentrations (8-12). Correspondingly, C2 domains of Syt-7 were found to be 400-fold more sensitive to Ca^2+^ than Syt-1 and have the slowest membrane dissociation kinetics among the synaptotagmins (13, 14). Both of the individual C2 domains from Syt-7 bind more tightly to membranes in the presence of Ca^2+^ than their respective counterparts in Syt-1 (15-18).

Ca^2+^-dependent membrane binding of synaptotagmin C2 domains is mediated by conserved aspartate residues in each domain’s Ca^2+^-binding loops (CBLs), of which CBL1 and CBL3 insert into membranes upon binding Ca^2+^ (19-25). The structural origins of membrane binding differences between Syt-1 and Syt-7 are not yet completely clear, although some features that contribute to the strong membrane binding of Syt-7 have been identified (16, 17, 26). In addition to their Ca^2+^-dependent membrane binding, the C2B domains of both isoforms contain a polybasic lysine-rich patch centered on the β-4 strand, which has been shown to bind preferentially to polyanionic phospholipids such as phosphatidylinositol-(4,5)-bisphosphate (PIP_2_) in a manner that is partially Ca^2+^-independent (17, 27-30). The stronger affinity of Syt-1 C2AB for membranes containing PIP_2_ manifests primarily through slower dissociation kinetics (*k*_off_), whereas association rates (*k*_on_) are essentially independent of lipid composition (30).

Mutational studies of Syt-1 and Syt-7 have revealed an interesting difference between the two proteins in the relative functional importance of the two C2 domains. The ability of Syt-1 to trigger fusion appears to depend more on Ca^2+^ binding by its C2B domain, whereas Syt-7 appears to be dominated by its C2A domain. In particular, neutralization of Ca^2+^-binding aspartate residues in the C2B domain of Syt-1 attenuates synchronous neurotransmitter release much more severely than corresponding mutations in the C2A domain (31, 32). However, fusion events mediated by Syt-7 display the opposite pattern: they are more sensitive to mutations in the Ca^2+^-coordinating residues of the Syt-7 C2A domain (9, 11, 33). In vitro membrane binding measurements also indicate that C2B dominates membrane binding in Syt-1 while C2A dominates in Syt-7 (17). In principle, differences between how these two sets of C2 domains work together could reflect functional specialization of the synaptotagmin isoforms.

In light of these findings, it has become essential to understand how the C2A and C2B domains work together to bind and insert into membranes, including comparisons between Syt-1 and Syt-7. Several studies have addressed this question for Syt-1, and have found that its C2 domains can either bind to the same target membrane or bridge between two opposing membrane surfaces (24, 34, 35). Indeed, a combination of these modalities can exist simultaneously depending on the protein used, the lipid composition, and the protein-to-lipid ratio (29, 36). It has also been reported that Syt-1 C2AB membrane binding is cooperative, on the basis of observations that both C2 domains co-penetrate a target membrane more deeply when present as the C2AB tandem than as isolated individual domains (22, 37). Biophysically, cooperativity implies that additional free energy stabilization is achieved by the combination of the C2A and C2B domains, beyond that which would be expected from simply linking any two membrane-binding protein domains together, i.e. from avidity effects (38). This cooperative model has been supported by single-molecule force measurements on Syt-1, for which C2AB domain dissociation from a target membrane was detected as single events that required a greater pulling force than the individual domains (39, 40). Additionally, a study replaced the C2A-C2B linker region of Syt-1 with a rigid polyproline helix and observed length-dependent effects on both membrane binding in vitro and secretion from neurons, further suggesting that co-insertion of the two domains may be functionally important (41).

In contrast to this model of cooperative membrane binding by the C2 domains of Syt-1, we have previously reported that the C2 domains of Syt-7 bind a target membrane independently (42). This assessment was made on the basis of both single-molecule diffusion measurements on planar supported lipid bilayers and dissociation kinetics from liposomes. In this model of independent membrane binding, the C2AB tandem domain is still predicted to have a stronger membrane affinity and Ca^2+^ sensitivity than the individual domains, due to avidity effects in which the binding strength of a multivalent protein-ligand interaction is enhanced due to the presence of multiple binding sites (38). However, the two binding sites do not strengthen one another’s affinity; i.e., two domains bind better than one but the effects are not synergistic.

In the present study, we sought to compare more directly the extent of cooperativity in C2AB membrane binding between Syt-1 and Syt-7. First, we measured Ca^2+^ sensitivity for Syt-7 C2A, C2B, and C2AB domains toward liposomes of physiological lipid compositions, with and without PIP_2_. Second, we performed direct side-by-side comparisons of liposome dissociation kinetics between Syt-1 and Syt-7 C2A, C2B, and C2AB domains, using the approach developed in our previous study (42). Finally, we applied an environment-sensitive fluorescent reporter assay with Syt-7 that was previously described and used to show co-penetration of the tandem C2 domains from Syt-1 (22). Together, the results show structural similarities but kinetic differences between C2AB domains of Syt-1 and Syt-7, which can be understood in terms of the relative energetics of membrane binding by these proteins’ component C2 domains.

## Materials and Methods

### Materials

1,2-Dioleoyl-sn-glycero-3-phosphocholine (DOPC), 1-palmitoyl-2-oleoyl-sn-glycero-3-phosphocholine (POPC), 1,2-dioleoyl-sn-glycero-3-phospho-L-serine (sodium salt) (DOPS), 1-palmitoyl-2-oleoyl-sn-glycero-3-phosphoserine (POPS), liver phosphatidylinositol (PI), 1-palmitoyl-2-oleoyl-sn-glycero-3-phosphoethanolamine (POPE), brain phosphatidylinositol-4,5-bisphosphate (PIP_2_), cholesterol (CH), and brain sphingomyelin (SM) were from Avanti Polar Lipids (Alabaster, AL). 1,2-dipalmitoyl-sn-glycero-3-phosphoethanolamine-N-(5-dimethylamino-1-naphthalenesulfonyl) (Dansyl-PE) was from NOF America (White Plains, NY). 3-[(3-Cholamidopropyl)dimethylammonio]-1-propanesulfonate (CHAPS) was from BioVision (Milpitas, CA). 5-({2-[(iodoacetyl)amino]ethyl}amino)naphthalene-1-sulfonic acid (IAEDANS) was from Molecular Probes (Eugene, OR). 2-Mercaptoethanol (βME) and ethylenediamine tetraacetic acid, tetrasodium salt (EDTA) were from Fisher Scientific (Hampton, NH) (≥ 99% purity). All reagents were American Chemical Society (ACS) grade or higher.

### Protein Cloning, Expression, and Purification

cDNA of human Syt-7 (IMAGE ID: 40121712) and Syt-1 (IMAGE ID: 6187902) were obtained from American Type Culture Collection. Sequences encoding the Syt-7 C2A domain (residues N135-S266), Syt-7 C2B domain (S261-A403), Syt-7 C2AB domain (N135-A403), Syt-1 C2A domain (K141-E272), Syt-1 C2B domain (S265-K422), and Syt-1 C2AB domain (K141-K422) were subcloned into a glutathione S-transferase (GST)-fusion vector developed previously (15, 43). All DNA sequences were verified using primer extension sequencing (Eton Bioscience, San Diego, CA). Plasmids were transformed into *Escherichia coli* BL-21 for protein expression.

All proteins were purified using glutathione affinity chromatography. Cells were lysed in lysis buffer (50 mM Tris, 400 mM NaCl, 1% triton X-100, 1 mM βME pH 7.5 with protease inhibitors) using a Sonics VibraCell sonicator with a 6-mm probe. For single domains, lysates were treated with DNase (2 U/mL) obtained from Sigma Aldrich (St. Louis, MO) for 30 minutes. Lysates were centrifuged to pellet insoluble matter, and supernatants were incubated with glutathione sepharose 4B beads (GE Healthcare, Chicago, IL) for 3 hours at 4 °C. After incubation, the beads were washed extensively with 50 mM Tris base, 400 mM NaCl, 1 mM βME, pH 7.5 and subsequently with 50 mM Tris, 1.1 M NaCl, 5 mM EDTA, 1 mM βME, pH 7.5. Beads were then exchanged into 50 mM Tris base, 150 mM NaCl, 0.05 mM EDTA, 1 mM βME pH 7.7 for cleavage with restriction grade thrombin (Novagen, Millipore Sigma, Billerica, MA), and eluted using the thrombin cleavage buffer or Buffer A (25 mM HEPES, 15 mM NaCl, 140 mM KCl, 0.5 mM MgCl_2_, pH 7.4) plus 1 mM βME. βME was omitted during purification of wild-type Syt-1 C2A. For Syt-7 C2AB, 1-3 mM CHAPS was added to all wash and elution buffers.

Proteins for Ca^2+^ dependence and kinetic experiments were immediately purified further via gel filtration (Syt-1 C2A) or cation exchange (all other C2 fragments) chromatography using an Akta Purifier FPLC system (GE Healthcare). Gel filtration was performed using a Superdex G75 10/300 GL column (GE Healthcare) in Buffer A. Cation exchange was performed using a HiTrap SP HP 5 mL column (GE Healthcare) in Buffer A plus 1 mM βME and protein was eluted with a gradient of NaCl. Representative chromatograms are shown in Figure S1. Protein integrity was confirmed using SDS-PAGE (Figure S2), and masses were further verified using MALDI. Absorbance spectra were measured using a Nanodrop 2000 spectrometer (Thermo Fisher Scientific) to assess removal of nucleic acids. All purified proteins had A_260_/A_280_ ratios ≤ 0.54, indicating the absence of nucleic acid contamination (44). Finally, protein concentrations were measured using UV absorbance (PerkinElmer) based on predicted extinction coefficients at 280 nm (http://protcalc.sourceforge.net). The purified proteins were aliquoted, flash-frozen, and stored at −80 °C. Prior to use, aliquots were thawed and centrifuged at 17000 ×g for 2 minutes to remove any debris, and the UV absorbance spectrum was re-checked to verify protein concentration and lack of nucleic acid.

### Liposome Preparation

Phospholipids in chloroform were combined at the desired molar ratio for each experiment (Table 1). After the evaporation of chloroform, the lipid films were dried under vacuum for ≥ 2 hours and rehydrated in Buffer A containing 10 mM βME to a final concentration of 3 mM total lipid. Small unilamellar vesicles (SUVs) were prepared by sonication to clarity on ice using a Sonics VibraCell sonicator with a 3-mm tip. Liposomes were stored at 4 °C for at least 8 hours after preparation, and were used within one week. Lipid concentrations are reported as total accessible lipid, which is assumed to be one-half of the total lipid present (i.e., lipids are assumed to be evenly distributed between inner and outer leaflets).

**Table 1.** Lipid compositions used in this study.

| Target Membrane Lipid Compositions (mol %) |  |  |  |  |  |  |  |  |
| --- | --- | --- | --- | --- | --- | --- | --- | --- |
| Name | PE | PC<sup>a</sup> | PS<sup>a</sup> | PI | PIP<sub>2</sub> | SM | CH | Dansyl-PE<sup>b</sup> |
| PM | 27.9 | 10.7 | 21.3 | 3.6<sup>c</sup> | 2.0<sup>c</sup> | 4.5 | 25 | 5.0 |
| PM(-)PIP<sub>2</sub> | 27.9 | 10.7 | 21.3 | 5.6 | - | 4.5 | 25 | 5.0 |
| 1:1 DOPC/DOPS | - | 47.5 | 47.5 | - | - | - | - | 5.0 |

<sup>a</sup>POPC and POPS were used for the physiological membranes, while DOPC and DOPS were used for the simplified lipid composition for consistency with prior experiments (42).
<sup>b</sup>For AEDANS fluorescence measurements, dansyl-PE was omitted and replaced with PC.
<sup>c</sup>For AEDANS fluorescence measurements, PM liposomes contained 4.6% PI and 1.0% PIP<sub>2</sub>.

### Equilibrium Measurement of Ca^2+^-Dependent Protein-to-Membrane FRET

Buffers were prepared using Chelex-treated Ca^2+^-free water. Liposomes were incubated with 10% (v/v) Chelex beads (Bio-Rad) overnight at 4 °C to remove residual Ca^2+^. Protein stocks were dialyzed into Ca^2+^-free Buffer A. Quartz cuvettes were soaked in 100 mM EDTA then rinsed extensively with Ca^2+^-free water prior to use. Steady-state fluorescence experiments were performed using a Photon Technology International QM-2000-6SE fluorescence spectrometer at 25 °C. Excitation slit width was 2.4 nm for Syt-7 C2B on PM liposomes, and 1 nm for all other samples; emission slit width was 8 nm. CaCl_2_ was titrated into an initially Ca^2+^-free solution containing protein (0.3 μM) and liposomes (75 μM accessible lipid). Due to the extreme Ca^2+^ sensitivity of Syt-7 C2 domains, a Ca^2+^ buffering system containing 1.5 mM nitrilotriacetic acid (NTA) was used in order to maintain total [Ca^2+^] in excess of protein, as described previously (15). Concentrations of free Ca^2+^ and Mg^2+^ (the latter held constant at 0.5 mM) were calculated using MaxChelator (http://maxchelator.stanford.edu). Fluorescence resonance energy transfer (FRET) was measured (λ_excitation_= 284 nm, λ_emission_ = 510 nm) over a 10-s integration time for each cuvette. Each intensity value was corrected for dilution, and the intensity of a blank sample containing only buffer, lipid, and Ca^2+^ was subtracted. Reversibility was demonstrated by adding excess EDTA after titrations (Figure S3). In order to determine the Ca_1/2_ and Hill coefficient of each C2 domain, normalized data were fitted to the Hill equation,

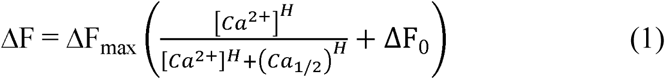

where ΔF is the fluorescence increase, H is the Hill coefficient, ΔF_0_ is the fluorescence change in the absence of Ca^2+^, and ΔF_max_ is the calculated maximal fluorescence change. Fitting was performed using Kaleidagraph 4.5 (Synergy Software). Data in figures are shown following normalization of ΔF_max_ to unity for each titration.

### Stopped-Flow Spectroscopy

Stopped-flow fluorescence kinetic measurements were performed using a BioLogic SFM3000 spectrophotometer (Knoxville, TN) using 284 nm excitation and a 455 nm long-pass emission filter. Unless otherwise noted, protein concentrations used were 1 μM for Syt-7 C2 domains and Syt-1 C2AB, and 5 μM for individual Syt-1 C2 domains (all concentrations listed are before mixing). Protein-to-membrane FRET (dansyl-PE emission) was monitored following rapid mixing of equal volumes of protein-bound liposomes (200 μM total accessible lipid, 200 μM CaCl_2_) and 2 mM EDTA in Buffer A. Dead time is estimated to be 1.4 ms. Data sets for each sample were calculated as the average of 8 or more time courses, and were fitted to a single-or double-exponential function (equation 2 or 3, respectively):

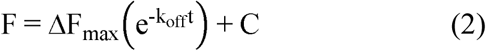

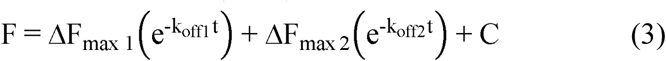

where the k_off_ are dissociation rate constants and C is an offset. C was subtracted and ΔF_max_ (or ΔF_max 1_ + ΔF_max 2_) normalized to unity in the figures shown. Rate constants listed are averages ± standard deviation of ≥ 3 independent replicate measurements.

### Dynamic Light Scattering

The *Z*-average diameter of liposome suspensions (1:1 DOPC/DOPS, 200 µM accessible lipids, 100 µM Ca^2+^) was determined using a Zetasizer Nano S90 (Malvern Instruments) before and after addition of protein. The samples were removed from the DLS instrument and used for stopped-flow kinetic measurements as described above.

### Purification of AEDANS-labeled Proteins

In order to facilitate labeling, native cysteine residues were removed from the proteins and unique cysteines were inserted using site-directed mutagenesis (Quik-Change II XL, Agilent). A Cys to Ala substitution was performed at C275 in Syt-7 C2AB and C2B. (Note: the C275A mutant of Syt-7 C2B was also used for all Syt-7 C2B kinetic experiments, as this mutation greatly simplified purification of the protein which otherwise had a propensity to form disulfide-linked dimers.) C260 was mutated to Ser in Syt-7 C2A and C2AB; we have previously reported that this mutation has only minor effects on the C2A domain (16). For AEDANS attachment, the following residues in the calcium binding loops 1 and 3 of the Syt-7 C2A, C2B, and C2AB domain were mutated to cysteine: T170 (C2A loop 1), N232 (C2A loop 3), T301 (C2B loop 1), and N364 (C2B loop 3). Mutations were confirmed using primer-extension sequencing (Eton Biosciences). Proteins were expressed and purified using glutathione affinity chromatography as described above, except that 0.5 mM TCEP was used in place of 1 mM βME for all domains, and 2.5 mM CaCl_2_ was included in the thrombin cleavage buffer for C2B. After thrombin cleavage and elution, protein concentrations were estimated from absorbance at 280 nm, and the proteins were incubated with IAEDANS (4:1 molar ratio of IAEDANS to protein). For single domains, proteins were labeled in 140 mM KCl, 0.5 mM MgCl_2_, 25 mM HEPES, 50 mM glutamic acid, 50 mM arginine pH 7.1 overnight at 4 °C (45). For C2AB tandems, proteins were labeled in thrombin cleavage buffer for 1 hour at room temperature. The reaction was quenched by adding 1 mM βME. To remove free dye and anionic contaminants, the proteins were further purified using cation exchange chromatography as described above, including 20 mM CaCl_2_ in the chromatography buffers. Following chromatography, proteins were exchanged into Buffer A plus 1 mM βME. Protein mass and labeling was verified using SDS-PAGE gel electrophoresis with visualization under ultraviolet light, and concentration was quantified based on AEDANS absorbance.

### AEDANS fluorescence assays

AEDANS fluorescence was measured using a Photon Technology International QM-2000-6SE fluorescence spectrometer at 25 °C. Fluorescence emission scans were measured scanning from 450 to 600 nm, with λ_excitation_ = 337 nm. Excitation slit widths were 1 nm except the following samples: C2A^3^B, 1.6 nm; C2AB^1^, 1.6 nm; C2AB^3^, 2.4 nm. Emission slit widths were 8 nm for all measurements. All measurements were carried out with 0.5 – 1 μM protein in Buffer A with 0.2 – 1.0 mM Ca^2+^ before and after addition of liposomes (33 μM total lipid).

## Results

### Strategy and protein purification

This study was designed to compare C2A-C2B interdomain cooperativity in Syt-1 and Syt-7; that is, whether a greater energetic stabilization is afforded by membrane binding for each C2AB tandem compared to the sum of the individual C2A and C2B domains. Thus, we expressed and purified individual domains and C2AB tandems of both Syt-1 and Syt-7 (Figures S1-S2). Initial purification was accomplished using affinity chromatography with cleavable GST tags. Because the Syt-1 C2B domain and both Syt-7 C2 domains are cationic and tend to co-purify with anionic contaminants, these individual domains and both C2AB tandem domains were additionally purified using cation exchange chromatography (Figure S1) (15, 17, 46). Syt-1 C2A was purified using glutathione affinity chromatography followed by gel filtration (Figure S1). Purity of all six protein domains was excellent as assessed using SDS-PAGE, UV absorbance, and MALDI mass spectrometry (see Methods).

### Ca^2+^ Sensitivity of Syt7 C2 Domains

The Ca^2+^ sensitivities and cooperativities of membrane binding were compared among individual and tandem C2 domains of Syt-7 (Figure 1, Table 2). For many C2 domains, including those from synaptotagmins, titration with Ca^2+^ in the presence of liposome membranes produces sigmoidal binding curves with a characteristic Ca_1/2_ ([Ca^2+^] at which the protein is half-maximally membrane-bound) and a Hill coefficient > 1 arising from cooperative binding of multiple Ca^2+^ ions and membrane to each domain (47). In order to compare these parameters for Syt-7 C2A, C2B, and C2AB domains, Ca^2+^ was titrated into solutions containing protein and liposomes with lipid compositions approximating the interior leaflet of the plasma membrane, with or without PIP_2_ (Table 1), and protein-to-membrane FRET was monitored (48). Due to the strong Ca^2+^ sensitivity of Syt-7, solutions were treated to remove as much free Ca^2+^ as possible and a Ca^2+^ buffering system was used to enable reliable measurement of binding even at free Ca^2+^ concentrations less than the protein concentration (15). Consistent with a previous report, the Syt-7 C2A domain bound liposomes containing a physiological membrane composition without PIP_2_ [PM(-)PIP_2_, Table 1] with a Ca_1/2_ around 5 μM and a Hill coefficient of 2.2 (Table 2) (15). Inclusion of PIP_2_ in the membrane had little impact on the Ca_1/2_ and Hill coefficient for this domain (Table 2).

**Figure 1.**
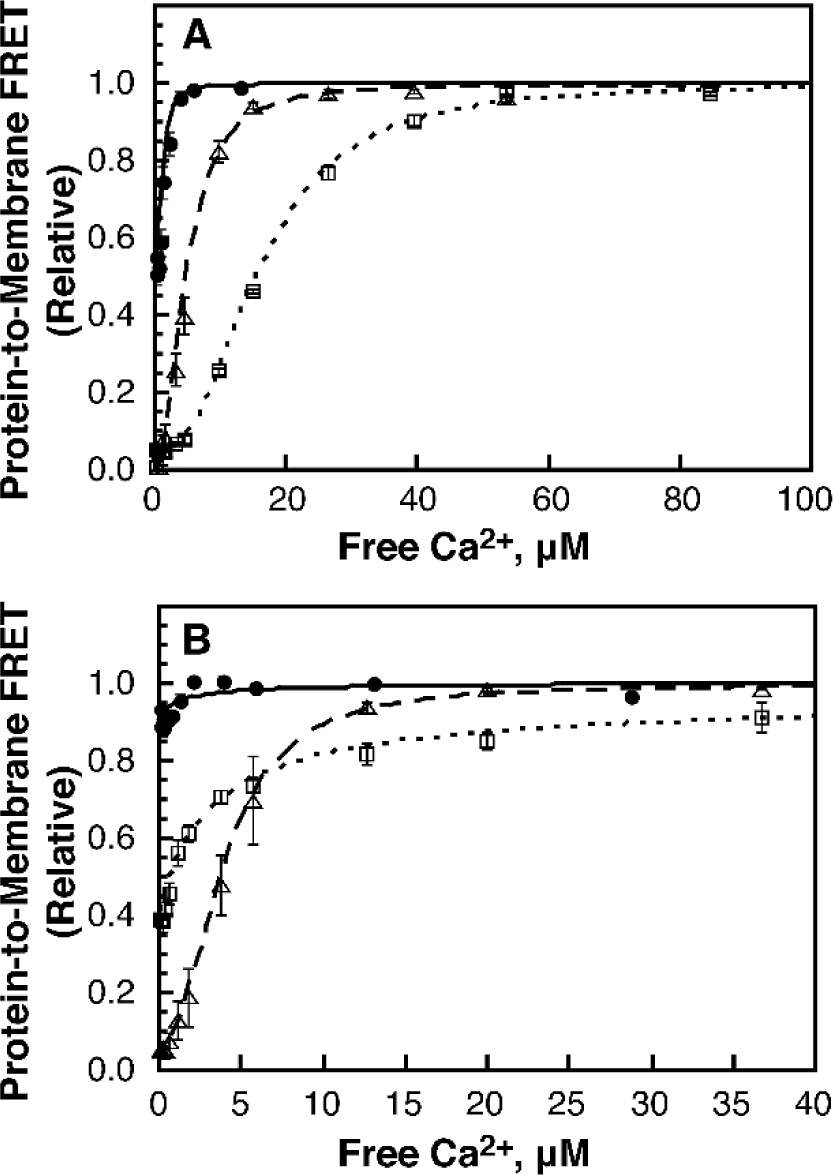
Ca^2+^ sensitivity of Syt-7 C2 domains. CaCl_2_ was titrated into solutions containing 0.5 μM Syt-7 C2A (open triangles, long-dashed curves), C2B (open squares, short-dashed curves), or C2AB (filled circles, solid curves) in the presence of (**A**) PM(-)PIP_2_ liposomes or (**B**) PM liposomes. Points and error bars shown are mean ± standard deviation of three independent replicate titrations; where not visible, error bars are smaller than the symbol. Data were normalized based on best fits to Eq. 1; the parameters from these fits are given in Table 2.

**Table 2.**
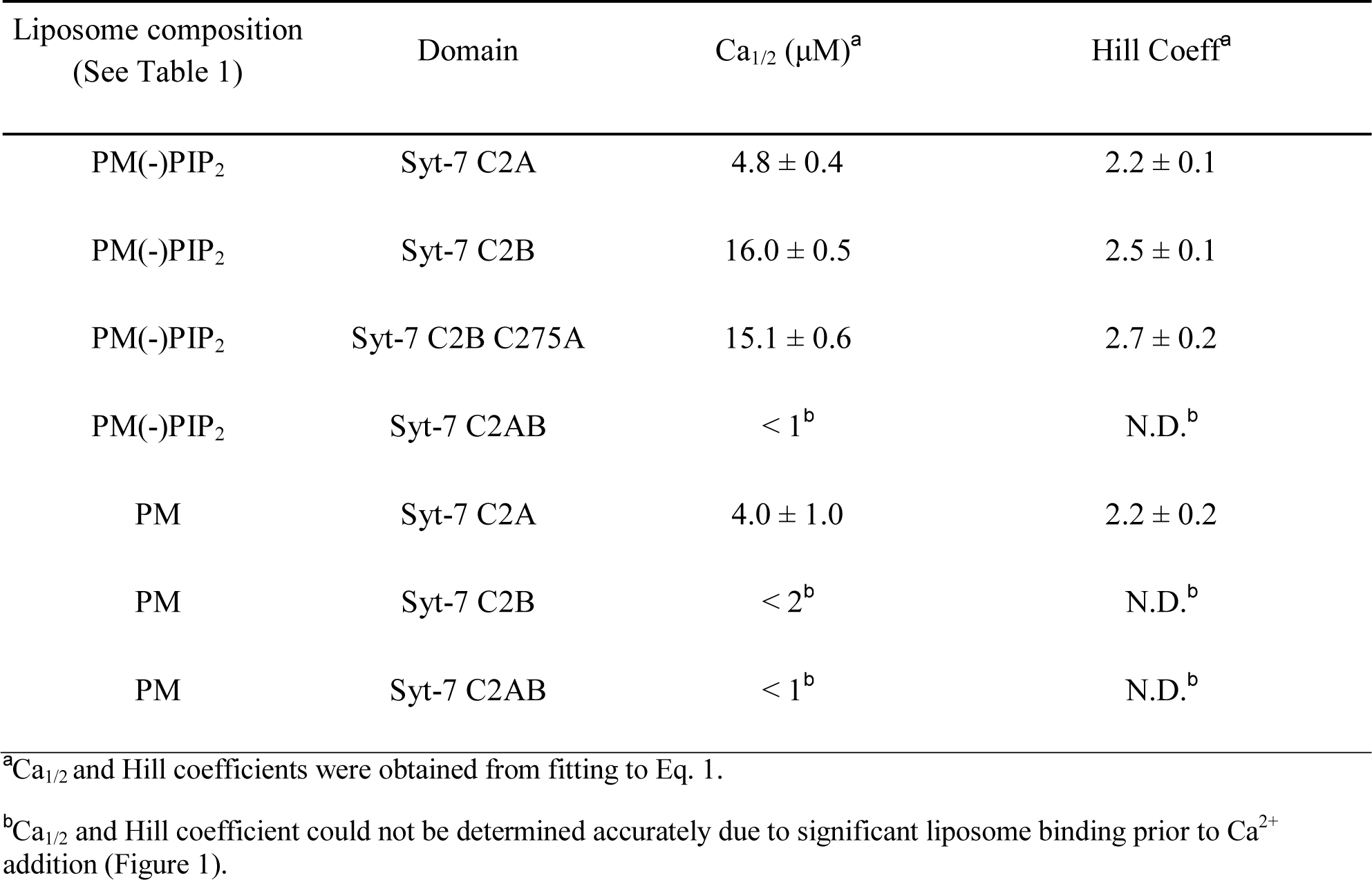
Equilibrium Ca^2+^ titration parameters of Syt-7 C2 domains

In contrast, the Syt-7 C2B domain showed a pronounced PIP_2_ dependence, consistent with a previous report (17). In the absence of PIP_2_, Syt-7 C2B bound liposome membranes with somewhat weaker Ca^2+^ sensitivity than the C2A domain (Figure 1A), but the trend was reversed upon inclusion of 1% PIP_2_ in the membrane, as the protein was partially membrane-bound even prior to Ca^2+^ addition (Figure 1B). This result suggests that Syt-7 C2B has a binding mode to PIP_2_ that is either Ca^2+^-insensitive or has a Ca_1/2_ near the limit of measurement. We estimate the level of Ca^2+^ contamination in these measurements to be < 1 μM; thus, the Syt-7 C2B domain can bind membranes containing PIP_2_ even at very low Ca^2+^ concentrations. This binding was reversed upon addition of the Ca^2+^ chelator EDTA (Figure S3). The Ca^2+^ dependence and Hill coefficient of the Syt7 C2B domain were not significantly affected by mutation of Cys275 to alanine (Table 2), a mutation which was required for the AEDANS measurements described below. Thus, all further experiments with Syt-7 C2B used the C275A variant.

The Syt-7 C2AB tandem had a lower Ca_1/2_ toward PM(-)PIP_2_ liposomes than either individual domain, and was approximately 50% bound to these liposomes even prior to Ca^2+^ addition (Figure 1A). Inclusion of 1% PIP_2_ further enhanced membrane binding, as the C2AB tandem was > 90% bound to PM liposomes prior to Ca^2+^ addition (Figure 1B). Binding of C2AB to PM(-)PIP_2_, but not PM, liposomes was reversed upon EDTA addition (Figure S3). Thus, the combination of the two domains increases the Ca^2+^ sensitivity of membrane binding, and may enhance a Ca^2+^-insensitive binding mode. However, it is not clear from these data whether such an enhancement arises from cooperativity or avidity effects.

### Assessing Cooperativity in Syt-1 and Syt-7 C2AB Domains from Dissociation Kinetics

In order to probe whether the energetics of Syt-1 and Syt-7 C2AB membrane binding are cooperative, we measured the kinetics of protein-membrane dissociation upon addition of the Ca^2+^ chelator EDTA. In this experiment, protein-to-membrane FRET decreases following the addition of EDTA to protein-liposome complexes. The EDTA coordinates Ca^2+^ ions as each C2 domain leaves the membrane surface, rendering each domain’s dissociation irreversible. Thus, if the two C2 domains bind and release from membranes independently, then the dissociation kinetic profile of the C2AB tandem should be closely similar to that of the rate-limiting domain. We have previously used this method to demonstrate that C2AB domains of Syt-7 dissociate independently from liposomes composed of 3:1 DOPC/DOPS (42). On the other hand, if the two C2 domains of a C2AB tandem bind membranes cooperatively, then C2AB dissociation is expected to be significantly slower than both of the individual domains.

Using this approach, the tandem C2 domains of Syt-1 are clearly observed to bind membranes cooperatively. Although the Syt-1 C2A domain dissociated from 3:1 DOPC/DOPS too quickly to measure, we quantified dissociation kinetics of Syt-1 C2A, C2B, and C2AB using 1:1 DOPC/DOPS (Figure 2A). The individual Syt-1 C2 domains’ dissociation time courses fit well to single exponential profiles, with the C2A rate constant similar to previous reports (15, 49). Dissociation of the C2B domain was slower than the C2A domain, consistent with its stronger membrane binding (17, 30, 40).

**Figure 2.**
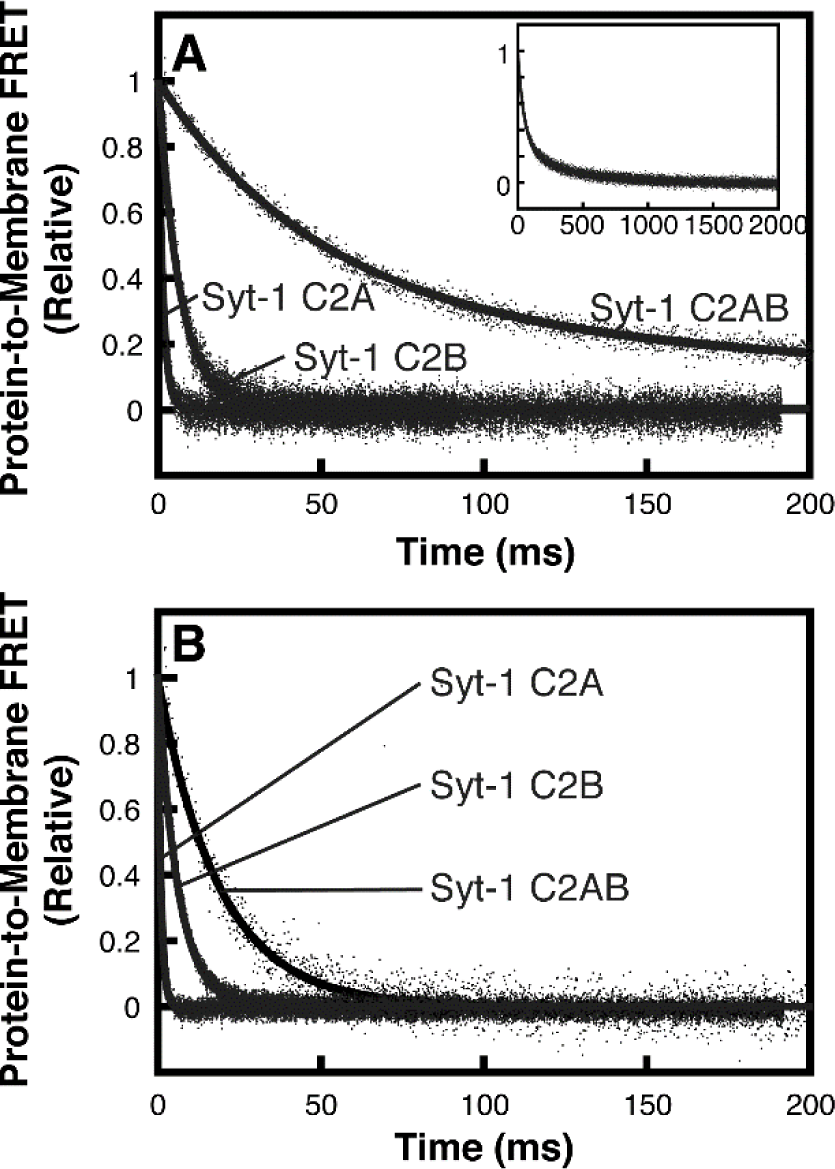
Dissociation kinetics of Syt-1 C2 domains on. (**A**) 1:1 DOPC/DOPS liposomes and (**B**) PM(-)PIP_2_ liposomes. Dansyl-PE fluorescence was monitored as solutions containing 5 µM (single domains) or 1 µM (tandem domain) protein, 200 µM CaCl_2_, and synthetic vesicles (200 µM accessible lipid) were rapidly mixed with an equal volume of 2 mM EDTA (all concentration listed are before mixing). Kinetic data were fit to Eq. 2 or Eq. 3, with rate constants shown in Table 3. Inset shows the full timescale of measurement of Syt-1 C2AB. Data shown are representative of ≥ 3 independent measurements.

Syt-1 C2AB dissociation was best fit by a double exponential, of which the faster rate constant was ∼7-fold slower than the isolated C2B domain and > 50-fold slower than the isolated C2A domain (Table 3). Similarly, Syt-1 C2AB dissociation from PM(-)PIP_2_ liposomes was at least 3-fold slower than either the C2A or C2B domain alone (Figure 2B, Table 3). Thus, the energy barrier for membrane release upon EDTA addition is significantly greater for the Syt-1 C2AB tandem than for both of its individual domains, consistent with a model of cooperative membrane insertion.

**Table 3.** Dissociation kinetics of Syt-1 and Syt-7 C2 domains. Double exponential rate constants are reported with their respective percent amplitudes from the best-fit curve.

| Domain | k<sub>off</sub>, 1:1 DOPC/DOPS (s<sup>-1</sup>) | k<sub>off</sub>, PM(-)PIP<sub>2</sub> (s<sup>-1</sup>) |
| --- | --- | --- |
| Syt-1 C2A | 710 ± 30 | 1020 ± 90 |
| Syt-1 C2B | 140 ± 10 | 170 ± 10 |
| Syt-1 C2AB | 21 ± 2 (72% amp) | 48 ± 5 |
|  | 2.6 ± 0.1 (28% amp) |  |
| Syt-7 C2A | 6.39 ± 0.04 | 7.4 ± 0.3 |
| Syt-7 C2B | 80 ± 9 (88% amp) | 100 ± 20 |
|  | 13 ± 2 (12% amp) |  |
| Syt-7 C2AB | 4.6 ± 0.3 (74% amp) | 5.9 ± 0.6 (77% amp) |
|  | 0.16 ± 0.04 (26% amp) | 0.64 ± 0.09 (23% amp) |

Dissociation kinetics of Syt-7 C2 domains are largely consistent with independent membrane binding, in contrast to Syt-1 (Figure 3). Dissociation of each individual Syt-7 domain was slower than that of the corresponding Syt-1 domain, consistent with previous reports (Table 3) (14, 15). The difference is particularly pronounced between the C2A domains, for which Syt-7 C2A dissociated >100-fold slower from both membrane compositions tested. Notably, for Syt-7 the C2A domain dissociated slower than the C2B domain, in contrast to Syt-1 for which the isolated C2B domain dissociates slower. This result is consistent with prior reports that the C2A domain dominates membrane binding and fusion properties for Syt-7 (9, 11, 17, 33).

**Figure 3.**
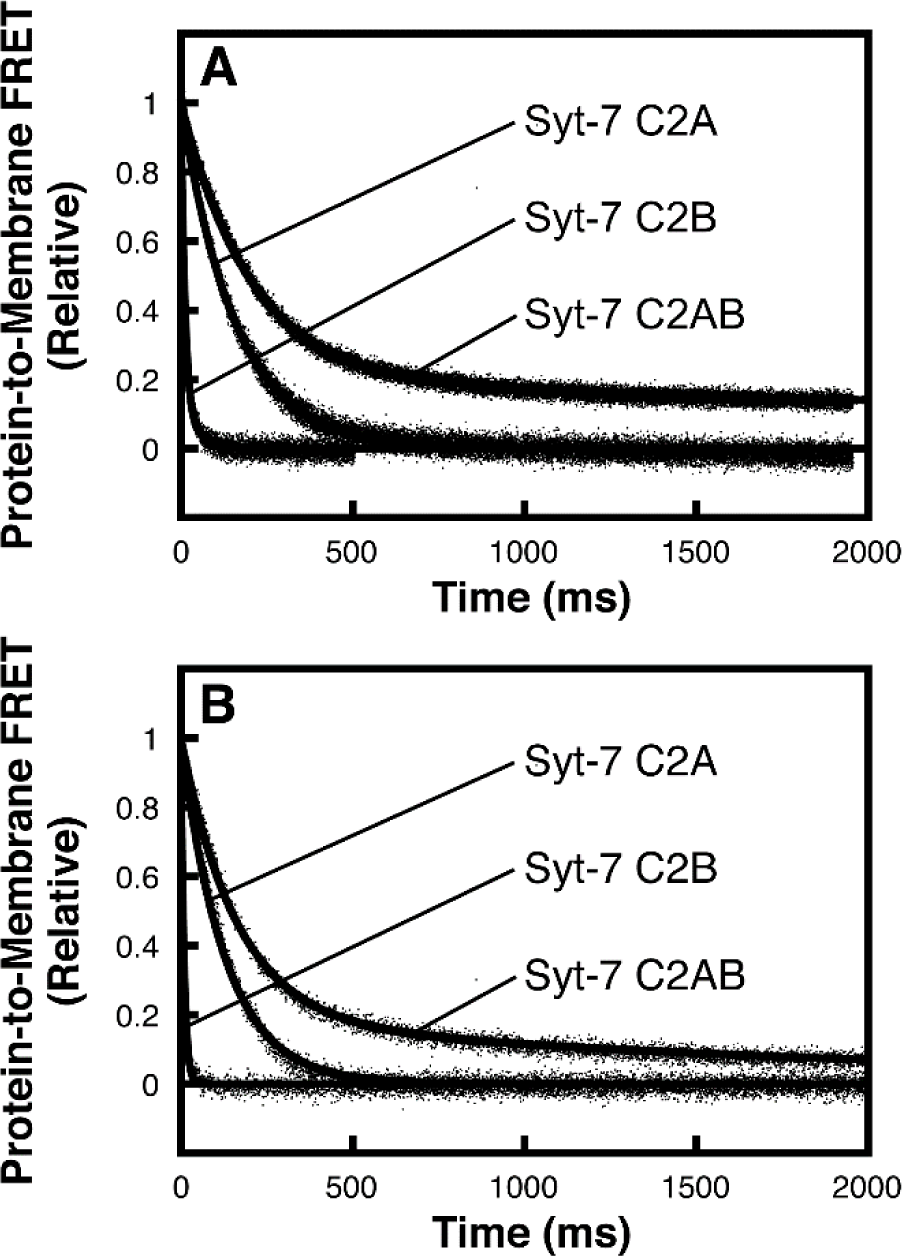
Dissociation kinetics of Syt-7 C2 domains on. (**A**) 1:1 DOPC/DOPS liposomes and (**B**) PM(-)PIP_2_ liposomes. Dansyl-PE fluorescence was monitored as solutions containing 1 µM protein (in panel A) or 0.25 µM protein (in panel B), 200 µM CaCl_2_, and liposomes (200 µM accessible lipid) were rapidly mixed with an equal volume of 2 mM EDTA (all concentrations listed are before mixing). Kinetic data were best fit to Eq. 2 or Eq. 3, with rate constants listed in Table 3. Data shown are representative of ≥ 3 independent measurements.

Dissociation of Syt-7 C2AB from both 1:1 DOPC/DOPS and PM(-)PIP_2_ was biphasic, with a major (faster) rate constant that differed by no more than 30% from the isolated C2A domain on each liposome composition tested (Figure 3, Table 3). The double exponential character suggests the presence of multiple populations, and one possibility is that a subpopulation (20-30%) of Syt-7 C2AB is bound to membranes in a cooperative fashion analogous to Syt-1 (see Discussion). However, the larger population of Syt-7 C2AB has dissociation kinetics that are more consistent with its two C2 domains binding membranes independently.

### Liposome Clustering by Syt-7 C2B

Some Syt C2 domains, including C2AB tandems, are known to promote clustering of liposomes in a Ca^2+^-dependent manner (17, 34, 35, 50, 51). This effect could conceivably result in slower dissociation rates, e.g. if a subpopulation of protein is initially inaccessible to the added EDTA. In order to probe whether liposome clustering occurs under the conditions used in the measurements above, we performed dynamic light scattering measurements of 1:1 DOPC/DOPS liposomes in the presence of different concentrations of Syt-7 C2B. Addition of 0.5 μM protein to liposomes resulted in a significant increase in particle size, from an average diameter of ∼100 nm to ∼500 nm (Figure S4A). Further addition of protein to 2 μM increased the size distribution further, to ∼800 nm (Figure S4B). These samples became visibly somewhat cloudy, but not flocculated, upon addition of 2 μM protein. Thus, Syt-7 C2B induces liposome clustering, but not large-scale aggregation, under conditions comparable to those used in the kinetic measurements reported above.

This degree of clustering appears to have no effect on the major dissociation rate constants. Kinetic profiles were measured using 0.5, 1, and 2 µM Syt7 C2B on the same concentration of liposomes (Figure S5). At the highest protein concentration, the dissociation profile was clearly double exponential, suggesting the presence of multiple populations. Kinetic profiles at the lower protein concentrations could also be fit to a double exponential, although the slower component was greatly decreased in intensity (Table S1). The slower phase was only a minor contributor (< 20%) to the overall kinetic profile at protein concentrations ≤ 1 μM. Importantly, the rate constants of the fast and slow components showed no trend among measurements using various concentrations of protein (Table S1). Thus, we conclude that liposome clustering and aggregation result in a subpopulation of protein with a significantly slower dissociation; however, at the protein concentrations used in the experiments above, the major, faster dissociation rate constant remains readily measurable. It is possible that liposome clustering may give rise to some of the minor, slower component(s) that we observe with Syt-1 and Syt-7 C2AB dissociation. However, these results support a high level of confidence that the major rate constants of the kinetic profiles correspond to straightforward protein-membrane dissociation that is not influenced by liposome aggregation effects.

### Membrane Insertion of Syt-7 C2 Domains

In order to test whether the tandem structure of Syt-7 C2AB affects its C2 domains’ membrane insertion, we used the environment-sensitive fluorescent probe AEDANS. This approach has been used previously to demonstrate membrane insertion of Syt-1 C2 domains (22, 23). AEDANS was attached to a unique engineered Cys residue on the tips of CBL1 and CBL3 on each Syt-7 C2 domain both individually and in the C2AB tandem (Figure S6). These are the regions that penetrate membranes in many C2 domains including those from synaptotagmins (16, 21, 52-55). Upon addition of physiological liposomes (PM(-)PIP2, Table 1) to protein in the presence of excess Ca^2+^, the AEDANS fluorescence emission increased and blue-shifted for both loops of C2A (Figure 4A-B) and loop 3 of C2B (Figure 5A-B). The increase in fluorescence emission was amplified for all four loops when measured in the C2AB tandem (Figures 4C-D and 5C-D). The difference between individual and tandem domains was especially striking for C2B loop 1, whose fluorescence increased dramatically upon liposome addition to the C2AB tandem but not the individual C2B domain (Figure 5A,C).

**Figure 4.**
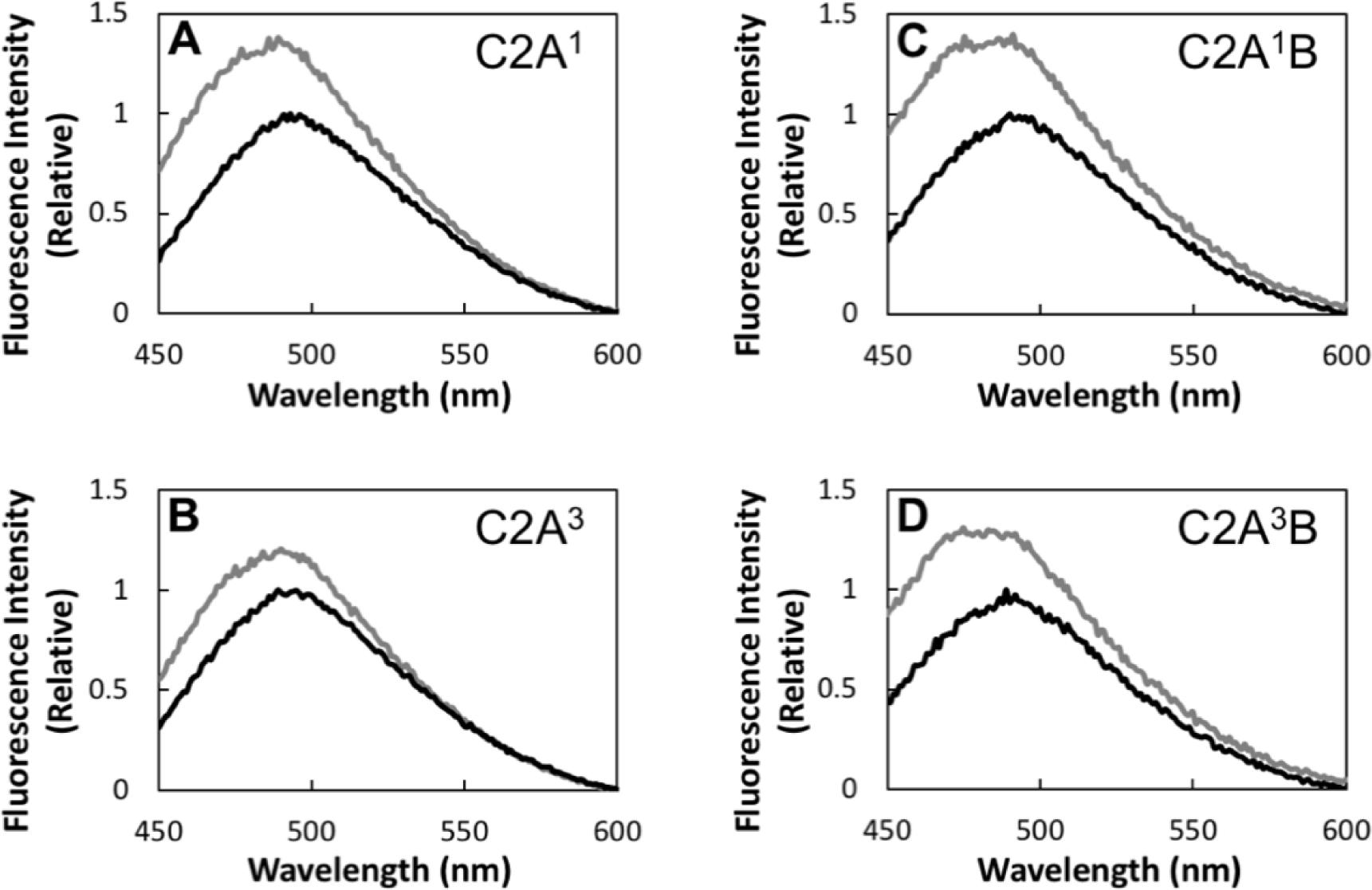
Membrane insertion of Syt-7 C2A domain loops in the absence of PIP_2_. Fluorescence emission spectra are shown of Syt-7 protein domains labeled with AEDANS at (**A**) C2A^1^, (**B)** C2A^3^, (**C**) C2A^1^B, or (**D**) C2A^3^B (see Figure S6 for schematic and Methods for labeled residues) in solution alone (black) and after addition of PM(-)PIP_2_ liposomes (gray). All spectra are normalized to the maximum intensity in the absence of lipid.

**Figure 5.**
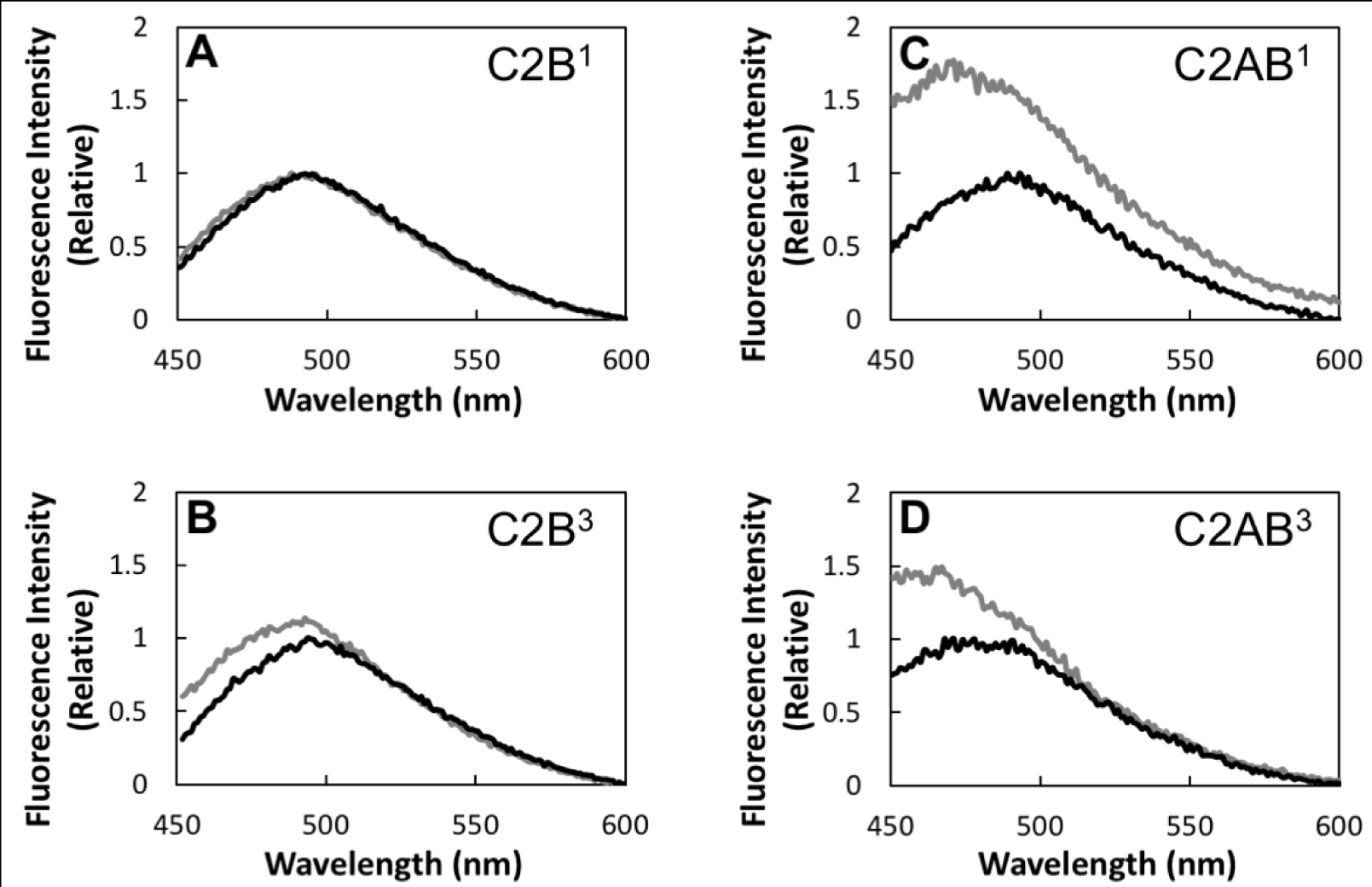
Membrane insertion of Syt-7 C2B domain loops in the absence of PIP_2_. Fluorescence emission spectra are shown of Syt-7 protein domains labeled with AEDANS at (**A**) C2B^1^, (**B**) C2B^3^, (**C**) C2AB^1^, or (**D**) C2AB^3^ (see Figure S6 for schematic and Methods for labeled residues) in solution alone (black) and after addition of PM(-)PIP_2_ liposomes (gray). All spectra are normalized to the maximum intensity in the absence of lipid.

Inclusion of 2% PIP_2_ in the liposome composition resulted in a similar pattern, except that the amount of fluorescence increase for the individual domains was somewhat greater than in the absence of PIP_2_ (Figures S7-S8). Overall, the C2 domains of Syt-7 penetrate into membranes when present as single or tandem domains. However, C2B in particular penetrates significantly more deeply when it is part of the C2AB tandem structure. This result is qualitatively similar to what was previously reported for Syt-1 (22).

## Discussion

In this study, we have employed FRET-based equilibrium Ca^2+^ titrations, stopped-flow kinetic measurements, and fluorescence-based membrane insertion assays to explore how the two C2 domains of Syt-7 work together in comparison to those of Syt-1. Four main conclusions can be drawn from the results: (i) the tandem C2 domains of Syt-1 bind membranes cooperatively, consistent with previous reports (22, 37); (ii) the major population of Syt-7 C2AB has dissociation kinetics that are consistent with independent, not cooperative, membrane binding (42); (iii) the Syt-7 C2AB domain is highly sensitive to Ca^2+^, even more so than its individual C2 domains, and (iv) the Ca^2+^-binding loops of Syt-7 C2B insert more deeply into membranes in the context of the C2AB tandem, similarly to Syt-1 (22); Although the last finding might appear to contradict a model of independent Syt-7 C2AB membrane binding, it can be understood in terms of relative contributions to binding energetics from the C2A and C2B domains, as discussed below.

### Cooperativity in Syt-1, Avidity in Syt-7

We previously reported that Syt-7 C2A and C2B domains bind independently to anionic lipid membranes, based on single-molecule lateral diffusion measurements and stopped-flow spectroscopy using 3:1 DOPC/DOPS membranes (42). For the present study, it was not possible to use the lateral diffusion approach with Syt-1 due to the fast dissociation of its C2A domain. Therefore, we measured stopped-flow kinetics of dissociation from 1:1 DOPC/DOPS and PM(-)PIP_2_ liposomes, two compositions for which dissociation rate constants were within the measurable range of the stopped-flow spectrometer for all of the protein domains investigated. These results show a clear distinction between the two proteins: for Syt-1, the C2AB tandem dissociates much slower than either of its constituent domains (Figure 2), whereas the largest population of Syt-7 C2AB dissociates at a rate comparable to its isolated C2A domain (Figure 3). All of the domains dissociated more slowly from 1:1 DOPC/DOPS than from PM(-)PIP_2_, likely due to the difference in anionic lipid content. Importantly, the trends among the various domains are generally the same for the two lipid compositions, except that a minor, slow phase was detected for Syt-1 C2AB and Syt-7 C2B on 1:1 DOPC/DOPS, possibly due to the higher anionic lipid content enabling additional liposome clustering via binding to secondary sites on the C2B domains. Thus, the present results support a model of independent membrane binding for the C2 domains of Syt-7, but cooperative binding for the C2 domains of Syt-1.

Dissociation of Syt-7 C2AB was biphasic on both lipid compositions tested, even at the low 0.25 μM protein concentration used with PM(-)PIP_2_. The minor, slower phase could arise from a subpopulation of protein that exhibits cooperative binding, a contribution from liposome aggregation, or a combination of these phenomena. Because the origin of heterogeneity in Syt-7 C2AB is not yet clear, we focus our analysis on the dominant population which has characteristics of independent membrane binding.

The AEDANS-based membrane insertion results with Syt-7 may shed light on the structural mechanism of cooperativity in Syt-1. Similarly to Syt-1, we show that the Ca^2+^-binding loops of Syt-7, especially CBL1 of the C2B domain, penetrate into membranes much more deeply as part of the C2AB tandem than in the isolated domains (14, 22). All four loops show robust insertion both with and without PIP_2_ in the context of the C2AB tandem (Figures 5C-D, 6C-D, S7C-D, and S8C-D). In particular, the CBL3 of C2B inserts much deeper in the C2AB tandem than in the isolated C2B domain (Figure 5B,D and Figure S8B,D). For Syt-1, the C2B domain binds membranes much more tightly than C2A, even in the absence of PIP_2_ (Figure 2). In contrast, Syt-7 C2A binds membranes more tightly than Syt-7 C2B, at least in the absence of PIP_2_ (Figure 1A; Figure 3) (17). Thus, both our present results and previously reported data are consistent with a model in which the C2B domain of both isoforms is more strongly impacted by the C2AB linkage (Figure 6) (22). We suggest that the deeper insertion of the C2B domain significantly slows the membrane dissociation kinetics of Syt-1 C2AB because that protein’s membrane binding strength is dominated by its C2B domain. For Syt-7, membrane binding is dominated by the C2A domain, which does not penetrate much deeper in the C2AB tandem. This model may explain how the energetic effects of C2A-C2B linkage differ between Syt-1 and Syt-7, even though its structural effects on membrane binding are similar for the two proteins.

**Figure 6.**
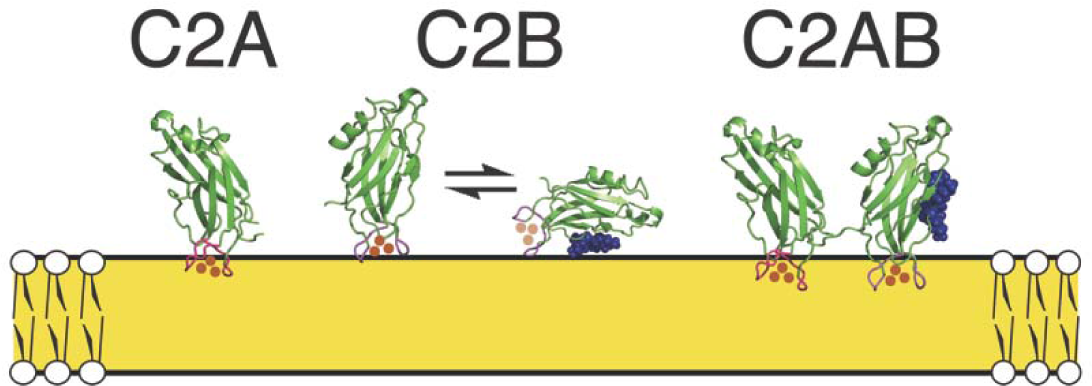
Structural model of linkage effects on insertion and dissociation kinetics. Syt-7 domains are drawn from published structures of C2A (PDB ID 2D8K) and C2B (56), although the model is consistent with previously reported structural data for Syt-1 (22, 30, 37). (*Left*) The isolated C2A domain penetrates membranes primarily via its CBLs (16, 54). (*Center*) The isolated C2B domain binds membranes relatively superficially through its CBLs and/or via interaction of its polybasic β-4 region (blue spheres) with lipid headgroups. (*Right*) In the C2AB tandem, both domains but especially C2B penetrate membranes more deeply. For Syt-1, because the C2B domain binds membranes relatively strongly (i.e., dissociates more slowly) in comparison to the C2A domain, its deeper insertion in the C2AB tandem leads to apparent interdomain cooperativity. For Syt-7, the isolated C2A domain binds membranes more tightly than the C2B domain, at least in the absence of PIP_2_. Because C2A insertion is only modestly affected by C2AB linkage, the deeper insertion of the C2AB tandem does not contribute significantly to binding energetics and the Syt-7 C2 domains do not display cooperative membrane binding.

### Apparent Ca^2+^ Independence

Syt-7 C2AB bound to membranes containing PIP_2_ even prior to Ca^2+^ addition (Figure 1B), and was not reversed by EDTA (Figure S3F). For this reason, we did not measure EDTA-induced dissociation kinetics from liposomes containing PIP_2_. Thus, it remains unknown whether Syt-7 membrane binding is cooperative in the presence of PIP_2_, although the Ca^2+^ and EDTA independence of the C2AB tandem suggests a strong effect of combining the two domains. In the presence of PIP, the membrane binding of the Syt-7 C2B domain becomes comparable to or stronger than the C2A domain (Figure 1B), therefore cooperativity under those conditions would be consistent with the model described above (Figure 6). Membrane bridging and liposome clustering could also contribute to the observed lack of EDTA reversibility for Syt-7 C2AB, but this effect is unlikely to be solely responsible because liposome clustering by other Syt C2 domains is reversed upon EDTA addition (50, 51).

The Syt-7 C2AB tandem was also partially membrane-bound prior to Ca^2+^ addition in the absence of PIP_2_; however, we interpret this result as arising from extreme sensitivity to sub-micromolar levels of Ca^2+^ in the assay solutions rather than Ca^2+^ insensitivity, as the C2AB tandem was effectively removed from PM(-)PIP_2_ liposomes upon EDTA addition (Figure S3C). Furthermore, neither individual domain of Syt-7 was observed to have a Ca^2+^-insensitive component in the absence of PIP_2_ (Figure 1A). The isolated Syt-7 C2B domain also bound somewhat to PIP_2_-containing membranes prior to Ca^2+^ addition (Figure 1B); this result is reminiscent of prior reports of partial Ca^2+^-independent PIP_2_ binding by Syt-1 C2B (28, 57). This interaction was also reversed by EDTA (Figure S3E), suggesting it may not be truly Ca^2+^ independent. However, we cannot exclude the possibility of a weak, Ca^2+^-independent interaction that becomes screened electrostatically by the EDTA polyanion (58).

### Liposome Clustering and Aggregation

Previously, Syt-1 C2B and C2AB as well as Syt-7 C2A and C2AB have been found to induce liposome aggregation (17, 34, 50, 51). In this study, we report that Syt-7 C2B shares the same property using 1:1 DOPC/DOPS liposomes. This contrasts with a previous report which observed little to no aggregation with Syt-7 C2B and 3:1 PC/PS liposomes (17). It is possible that the increased anionic content in our liposomes is sufficient to engage a second binding site in this domain, similar to Syt-1 C2B. Both of the polybasic regions that have been suggested to impact bilayer bridging by Syt-1 C2B are present in Syt-7 C2B as well (7, 29, 59). As expected, liposome clustering is concentration dependent (Figure S4) (28, 57). In these experiments, liposome clustering arising from bilayer bridging may reflect a separate structural state from the co-insertion events whose extent of cooperativity was probed in this study, and therefore we sought to minimize aggregation by choosing conditions that favor co-insertion, such as low protein-to-lipid ratios (36). At higher protein-to-lipid ratios, liposome aggregation appeared to produce or exaggerate a slow component of the kinetic time course (Figure S5; Table S1). However, the rate constant of the major component of the kinetic time course remained unaffected; therefore, we conclude that this rate constant represents dissociation of the proteins from the liposome surface.

### Implications for Evolution of Synaptotagmin Function

Despite their different Ca^2+^ sensitivities and kinetics, both Syt-1 and Syt-7 insert their C2B domain Ca^2+^-binding loops more deeply into membranes as C2AB tandems than as individual domains. Our data suggest this insertion leads to interdomain cooperativity in Syt-1 but not Syt-7, at least in the absence of PIP_2_, presumably because the enhanced C2B insertion increases overall membrane binding affinity more strongly for the C2B-dominated Syt-1 than for the C2A-dominated Syt-7 (Figure 6). The conservation of this structural feature suggests that co-insertion of C2 domain Ca^2+^ binding loops into the same membrane may be a common feature among synaptotagmins, even under conditions where some degree of liposome clustering is also present (36). We recently posited that synaptotagmins in the three-dimensional context of a fusion pore could assume a geometry in which the C2A and C2B Ca^2+^-binding loops co-penetrate the fusion pore neck while the polybasic regions bridge between opposing membranes (7). In other words, cooperative membrane insertion and bilayer bridging are not necessarily contradictory functions. In principle, cooperativity between C2 domains could facilitate rapid and efficient co-insertion to stabilize the fusion pore during a Ca^2+^ signaling event, while maintaining a low membrane affinity at basal Ca^2+^ levels. Future studies comparing structural and energetic properties of synaptotagmin isoforms could provide key insights into the evolution of synaptotagmin function in fusion pore formation and stabilization.

## Author Contributions

H.T., L.A., and J.K. designed research; H.T. and L.A. performed research; H.T., L.A., and J.K. analyzed data; H.T. and J.K. wrote the manuscript.

